# A live-cell platform to isolate phenotypically defined subpopulations for spatial multi-omic profiling

**DOI:** 10.1101/2023.02.28.530493

**Authors:** Tala O. Khatib, Angelica M. Amanso, Brian Pedro, Christina M. Knippler, Emily R. Summerbell, Najdat M. Zohbi, Jessica M. Konen, Janna K. Mouw, Adam I. Marcus

## Abstract

Numerous techniques have been employed to deconstruct the heterogeneity observed in normal and diseased cellular populations, including single cell RNA sequencing, *in situ* hybridization, and flow cytometry. While these approaches have revolutionized our understanding of heterogeneity, in isolation they cannot correlate phenotypic information within a physiologically relevant live-cell state, with molecular profiles. This inability to integrate a historical live-cell phenotype, such as invasiveness, cell:cell interactions, and changes in spatial positioning, with multi-omic data, creates a gap in understanding cellular heterogeneity. We sought to address this gap by employing lab technologies to design a detailed protocol, termed Spatiotemporal Genomics and Cellular Analysis (SaGA), for the precise imaging-based selection, isolation, and expansion of phenotypically distinct live-cells. We begin with cells stably expressing a photoconvertible fluorescent protein and employ live cell confocal microscopy to photoconvert a user-defined single cell or set of cells displaying a phenotype of interest. The total population is then extracted from its microenvironment, and the optically highlighted cells are isolated using fluorescence activated cell sorting. SaGA-isolated cells can then be subjected to multi-omics analysis or cellular propagation for *in vitro* or *in vivo* studies. This protocol can be applied to a variety of conditions, creating protocol flexibility for user-specific research interests. The SaGA technique can be accomplished in one workday by non-specialists and results in a phenotypically defined cellular subpopulation for integration with multi-omics techniques. We envision this approach providing multi-dimensional datasets exploring the relationship between live-cell phenotype and multi-omic heterogeneity within normal and diseased cellular populations.

## INTRODUCTION

Cellular heterogeneity underlies all biological systems. Heterogeneity exists across the varying stages of development, differentiation, and disease over length scales from DNA to organism^1–3^. This cellular heterogeneity emerges as a result of epigenetic, transcriptional, and post-translational diversity within and between populations^4,5^. These heterogeneous populations cooperate to maintain biological homeostasis, often providing a selective advantage by enabling a heightened response to stimuli, microenvironment, or selective pressures^6–8^. Additionally, pathological states emerge and progress under the influence of vast heterogeneity providing diseases, such as cancer, a myriad of potential mechanisms for therapeutic evasion, escape of immune surveillance, and relapse^9–11^. Ultimately, an effective understanding of the temporal progression for any normal or diseased biological system requires consideration of the interplay and cooperation between genetically, epigenetically, and phenotypically diverse cellular subpopulations.

Heterogeneous cellular populations orchestrate diverse phenotypic responses that can be imaged over space and time. However, technologies capable of deriving multi-omic analysis from live, spatiotemporally defined cellular populations are limited. Here, we address this gap in technology by applying live-cell microscopy to explore phenotypic cellular diversity and identify distinct subpopulations across cellular landscapes. This protocol takes a phenotype-driven, live-cell imaging approach to link historical cellular behavior with multi-omic and molecular information. We describe in detail how to exploit accessible lab technologies to isolate user-defined live cells with minimal space and time limitations.

### DEVELOPMENT OF THE PROTOCOL

Global multi*-*omics approaches are commonly utilized to test biological hypotheses, where bulk -*omics* techniques (*e.g.,* proteomics via mass spectrometry, RNA-and DNA-sequencing) are readily available and cost-effective^12,13^. For example, these technologies have driven critical discoveries in cancer research including insight into the regulation of onco-genes and proteins^14–16^. One notable drawback to “homogenizing” bulk multi*-* omics approaches is the inability to discern contributions from heterogeneous and rare subpopulations within each sample. More recent advances in single cell multi-omics provide insight toward resolving the distinct landscapes of subpopulations within a single population of cells; however, these approaches typically do not integrate phenotypic information within a physiologically relevant, live-cell state. Similarly, spatial multi-omics is a powerful tool to collect detailed molecular characterization of tissue while preserving spatial context, however samples are fixed and therefore cannot be propagated for further analysis^17^.

Tumor subpopulations incur distinct genomic and epigenetic profiles through selective pressures, increased genomic instability, and various degrees of entropy; these distinct subpopulations may drive unique invasive potentials and proliferative capacities, where cellular subpopulations cooperate to drive efficacious tumor progression and metastatic disease^11,13,18–25^. Despite our knowledge of this cellular heterogeneity, the mechanisms underlying pack formation, function, and impact on cancer progression are largely unknown. To probe subpopulation heterogeneity and its role in collective invasion, we sought to develop an image-guided technique that allows for precise *in situ* selection, isolation, and expansion of phenotypically distinct, live cells and populations during 3D invasion^26^. By combining 3D cell culture with accessible lab technologies – including live-cell confocal microscopy and fluorescence-activated cell sorting (FACS) – the <u>S</u>p<u>a</u>tiotemporal <u>G</u>enomic and Cellular <u>A</u>nalysis (SaGA) technique allows for the optical marking of single live cells (or collections of cells) within a defined region of interest using a photoconvertible tag, Dendra2 (Fig. 1). After photoconversion, the cells can be further observed *in situ* or removed from their environment and flow-sorted for a myriad of downstream analyses including, genotypic and phenotypic stability and/or flexibility of clones and subpopulations (Table 1). Alternatively, these extracted cells can be immediately processed for single or bulk cell multi-omics applications.

**Fig. 1.**
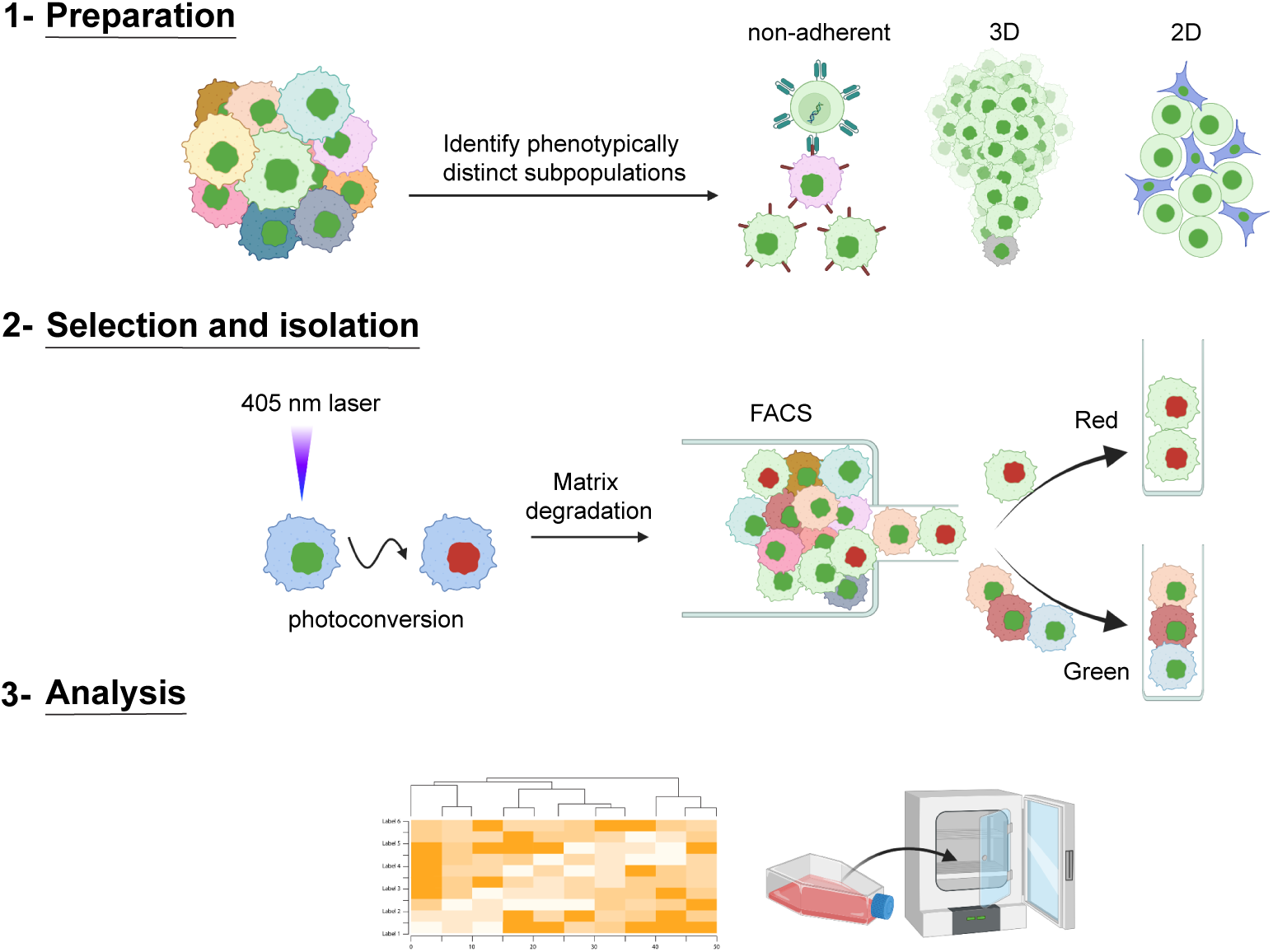
I SaGA schematic to isolate distinct cell(s) based upon live, user-defined phenotypic criteria. Schematic showing three broad steps of SaGA: 1) Preparation, 2) Selection and Isolation, and 3) Analysis. SaGA can be applied to a variety of cell conditions, such as non-adherent, 3-dimensional (3D), and 2-dimensional (2D), for selection, isolation, and analysis of live subpopulations within a parental population. Cells stably expressing a photoconvertible tag can be precisely photoconverted (from green to red) based upon live, user-defined, phenotypic criteria. These red photoconverted cells are then isolated utilizing fluorescence activated cell sorting (FACS) for multi-omic analysis and/or cell cultivation for long-term *in vitro* and *in vivo* analyses. Created with Biorender.com.

**Table 1.**
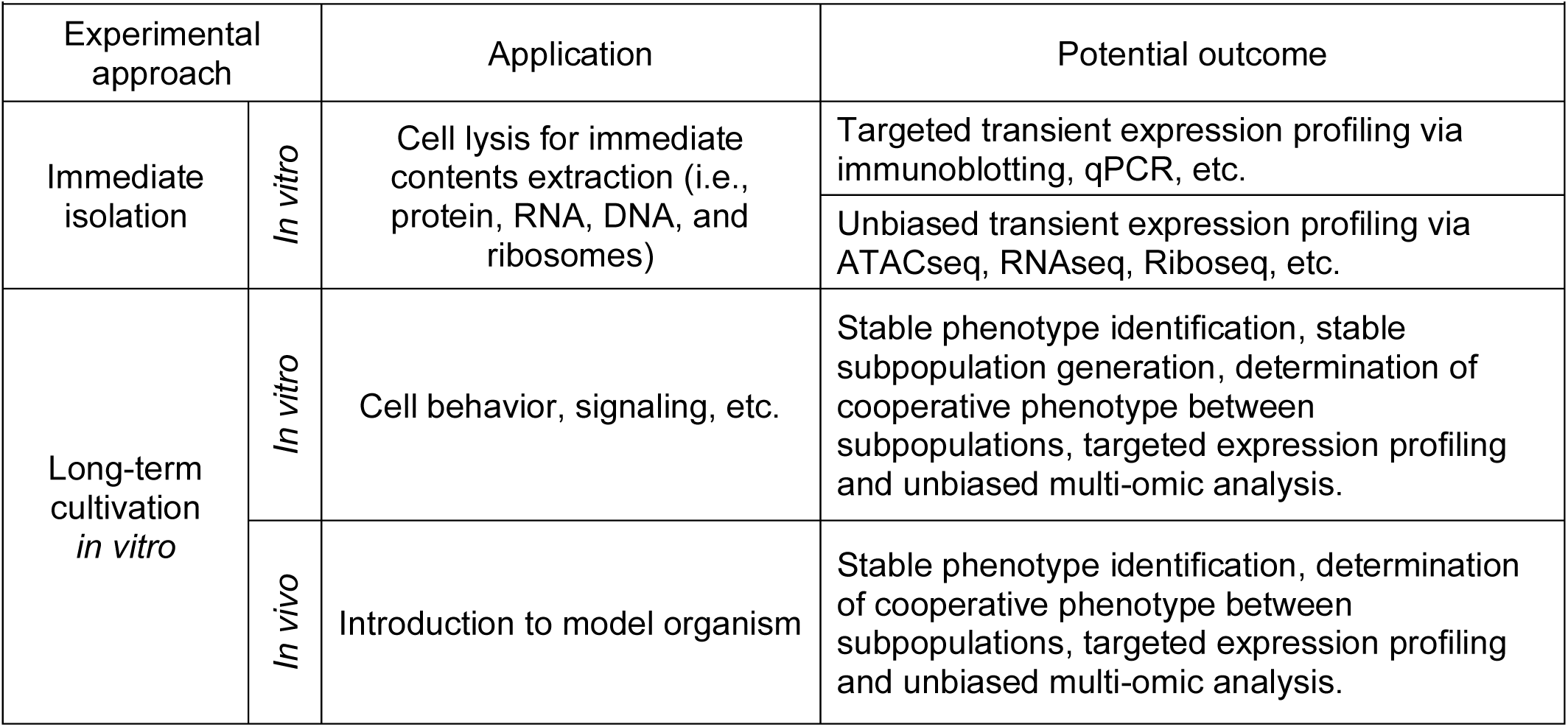
I Example downstream applications of SaGA-isolated subpopulations

Combining SaGA and multiple non-small cell lung cancer (NSCLC) lines, cell(s) were photoconverted based upon their spatial positioning within the collectively invading pack to isolate leader (front of the pack) and follower (trailing behind leaders) subpopulations^26^. To assess phenotypic stability overtime, cells were sorted for propagation and long-term phenotypic and transcriptomic analysis^26–30^. Epigenetic differences were assessed by performing a DNA methylation array, where cells were sorted for immediate DNA extraction and epigenetic analysis^26,28^. Taken together, these multi-omic results corroborate leader and follower spatial localization, providing, for the first time, a detailed mechanistic understanding of cellular positioning within the collective invasion pack.

### APPLICATION OF SAGA

The SaGA approach integrates standard laboratory practices to ask fundamental and clinically relevant questions about the mechanistic underpinnings of population heterogeneity. Prior to performing live-cell imaging, SaGA is flexible in experimental design and, therefore, adaptable toward a multitude of research-specific interests. Researchers can readily adopt this protocol with limited experience in confocal imaging or flow cytometry and may find applications in a range of fields including neuroscience or developmental biology^31–33^.

Using the H1299 NSCLC line, we found that leaders and followers isolated during 3D collective invasion have distinct genotypic, epigenetic, and phenotypic differences, and are phenotypically stable over many passages^26–30^. By mapping our bulk RNA sequencing results to the human reference genome Hg19 (GRCh37) and various filtering steps, we identified 14 distinct missense mutations between leaders and followers^30^. Similarly, epigenetic analysis via DNA methylation array featured global epigenetic rewiring in leaders compared to followers^28^. These results indicate that our NSCLC spatial localization is a coordinated patterning driven by genomic and epigenetic cellular profiles. At the RNA and protein levels, we found that underlying differences in their filopodia dynamics driven by a Jag1-Myo10 signaling axis further contribute to the stark differences in invasive capacities of the leaders and followers^28–30^. Similarly, invasive chains harbor metabolic heterogeneity, in which trailing followers are highly glycolytic and leaders depend upon mitochondrial respiration^27^. Taken together, these data highlight the ability to use phenotypic heterogeneity to decipher the genomic, epigenetic, and phenotypic underpinnings of tumor cell heterogeneity, and support the application of SaGA to investigate population heterogeneity.

Beyond phenotypic positioning within a collective invasion pack in NSCLC, SaGA can be applied to any image-able phenotype for selective enrichment. Applications include selection of cells based upon the sub-cellular localization of a protein of interest, proliferation rates, drug resistance, homo-or hetero-typic cellular interactions, and morphological changes due to differential cellular environments. During tissue and embryonic morphogenesis, complex architectural and temporal patterns of protein expression emerge. In this instance, SaGA can be readily utilized to answer questions governing cell fate decision making. Similarly, neurological diseases also highlight intercellular heterogeneity^34^. One example is Alzheimer’s disease in which microglia, specialized tissue-resident macrophages in the central nervous system, have distinct localization patterning^35^. In sum, SaGA provides a powerful platform for deconstructing live-cell phenotypic heterogeneity within any image-able, heterogenous cell population.

### EXPERIMENTAL DESIGN AND LIMITATIONS

Here, we elucidate key steps in the experimental strategy of SaGA, a method to isolate and evaluate the molecular dependencies of any image-able, phenotypically distinct cell subpopulation in live cell microscopy. Broadly, implementing SaGA requires a tissue culture grade facility, a laser scanning confocal microscope with a photoconversion regime, molecular biology approaches suitable for genetic manipulations, and a fluorescence activated cell sorter (FACS). Overall, the protocol includes stable cell transduction with a photoconvertible tag, live cell imaging to photoconvert the region of interest (ROI), FACS to isolate the cell subpopulation of interest, and downstream multi-dimensional analysis (Table 1, Fig. 2). These techniques are accompanied by a set of technical limitations, highlighting the importance of minimizing experimental bias and maintaining initial population heterogeneity (Fig. 3). In the following section, we discuss how to best accommodate these limitations to facilitate and maintain population heterogeneity (Fig. 3).

**Fig. 2.**
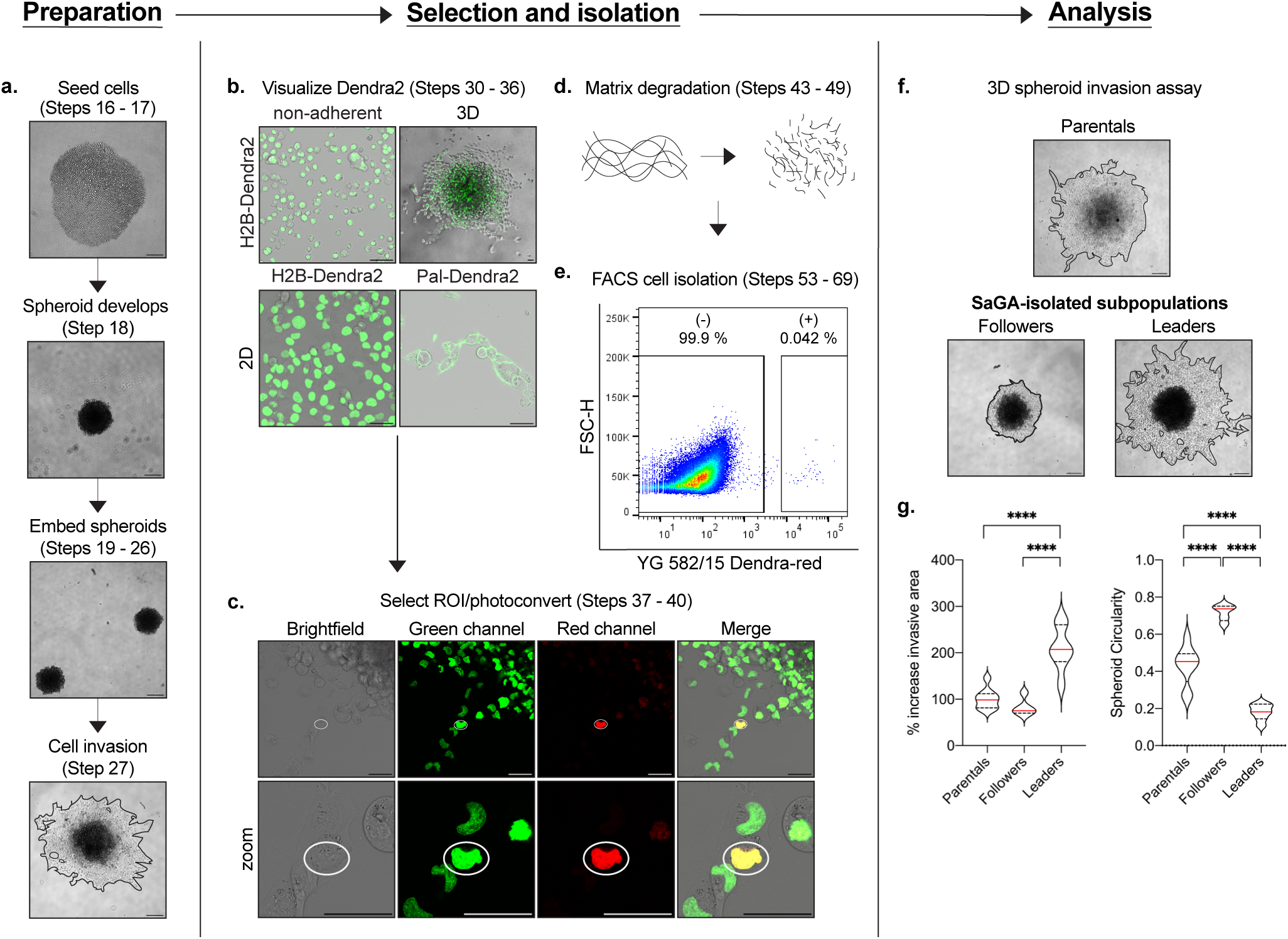
I SaGA workflow. Each panel provides an example of a major component of SaGA: preparation, selection and isolation, and analysis. **a.** 3D spheroid invasion assay set-up beginning with spheroid formation in a low adherence 96-well plate to embedment and invasion in recombinant basement membrane. Scale bar, 250 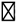m. **b.** Dendra2 visualization under non-adherent, 3D and 2D conditions. 2D conditions are shown utilizing both nuclear-(H2B-Dendra2) and membrane-(Pal-Dendra2) localized protein tags. Scale bar, 50 μm. **c.** Defining a region of interest (ROI) (white 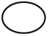) for cell selection and photoconversion. Scale bar, 50 μm. **d.** Matrix degrada-tion in 3D conditions utilizing collagenase/dispase cocktail. **e.** FACS plot showing non-photoconverted (-) and photoconverted (+) cells. **f.** 3D spheroid invasion assay with H1299 parental population and SaGA-isolated leader and follower subpopulations. Scale bar, 250 μm. **g.** Invasive area and spheroid circularity quantification. *p < 0.05 by one-way ANOVA with Tukey’s multiple comparisons test.

**Fig. 3.**
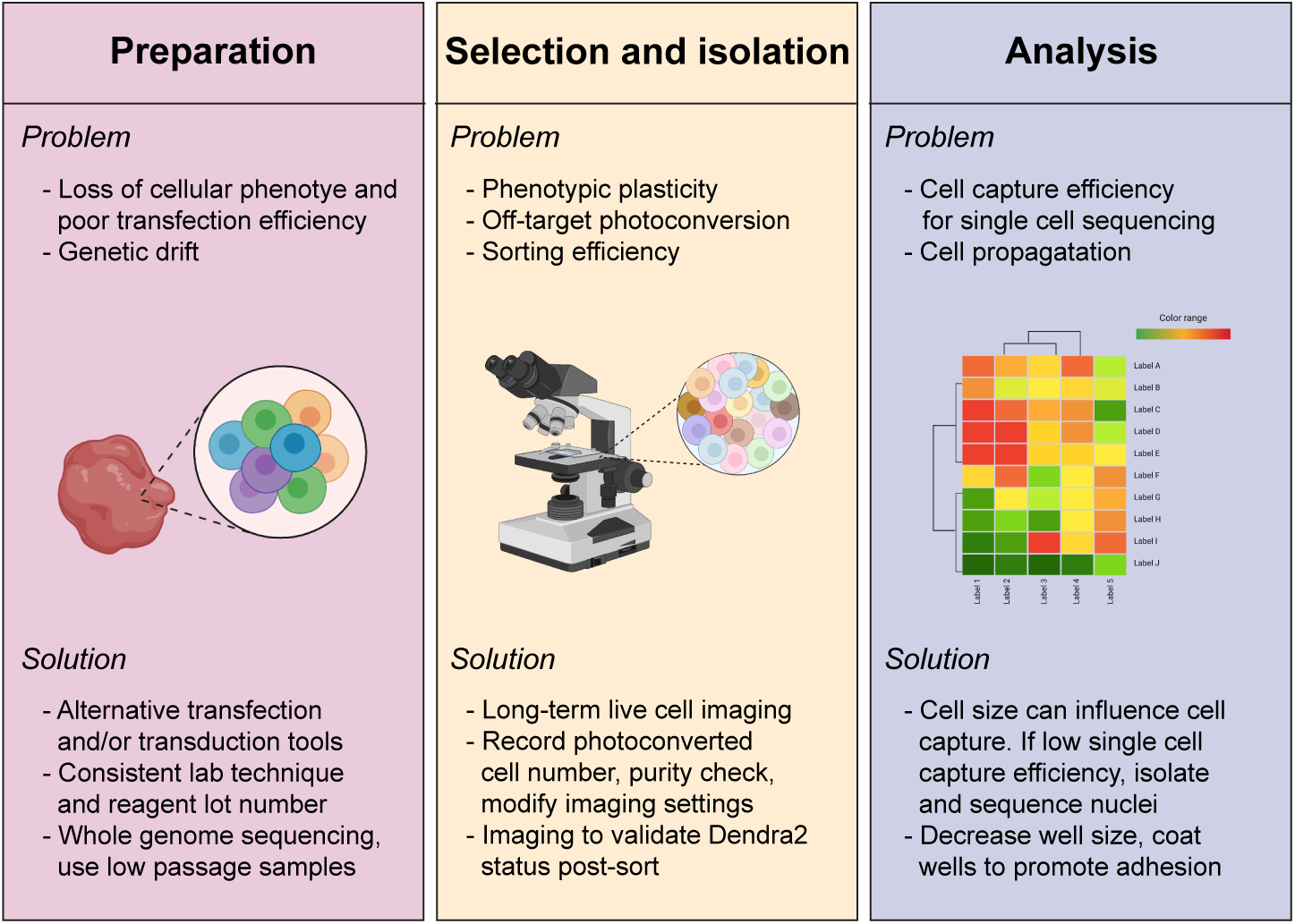
I Potential loss of heterogeneity and error sources and measures to minimize them. Cellular loss of heterogeneity can occur during sample preparation, selection and isolation, and analysis. Listed is each major stage of SaGA (preparation, selection and isolation, and analysis) with potential problems (bulleted above within each panel) that can occur and respective potential solutions (bulleted below within each panel). Graphical images created with Biorender.com.

#### Choice of fluorescent protein

Fluorescent tags have evolved over decades, from green fluorescent protein (GFP), discovered in 1962, to photoconvertible fluorescent proteins (PCFPs), discovered in 2002; PCFPs are characterized by their ability to switch emission spectra upon illumination with light at a specific wavelength and intensity, thereby allowing precise optical labeling and tracking of protein or cell dynamics^36–38^. Many PCFPs have been designed, including the green-to-red Dendra2 and mEos2 proteins, the orange-to-far-red PSmOrange protein, and the cyan-to-green PS-CFP2 protein^39–43^. These fluorescent molecules can also be targeted to distinct cellular regions through additional sequence modifications, allowing for precise photoconversion of sub-cellular structures such as the plasma membrane, nuclei, or mitochondria.

Choice of PCFP requires careful consideration of the experimental question, model and PCFP dynamics, and potential limitations. Previously published studies in our lab utilized SaGA to isolate cells based upon their location within a collective invasion pack. As such, to better distinguish the physical positioning between and amongst cells, we chose to use either a histone H2B-tagged Dendra2 (which localizes to the nucleus) or a palmitoylated Dendra2 (which localizes to lipid rafts, including those in the plasma membrane) for our experimentation. The Dendra2 PCFP is an engineered Kaede-like fluorescent monomeric protein with a light-driven covalent modification that results in an irreversible photoconversion^44,45^. Dendra2 can be initially excited at 490 nm to fluoresce in the GFP-like green fluorescent state (emission peak at 507 nm). Upon user-defined exposure to UV-violet or blue light, Dendra2 irreversibly photoconverts to a red fluorescent state (excitation/emission peaks at 553/573 nm)^36,38,46^. This PCFP has flexibility in that photoconversion can occur with either UV-violet (360 – 420 nm) or blue (460 – 500 nm) light excitement.

#### Cell transfection and transduction

Introducing genetic material into a cell can be performed stably or transiently with a myriad of well established biological, chemical, and physical methods^47–49^. While definitions vary in the literature, for clarity and the purposes of this protocol, we define transfection as the introduction of genetic material into a cell via non-viral methods, and transduction as the introduction via viral methods. The technique chosen depends upon the cell model and the experimental requirements. Ensuring adequate brightness and expression level and relatively homogenous PCFP expression is a key first step in an experimental design centering around maintaining representative phenotypic heterogeneity.

Our laboratory has used multiple techniques for stably introducing Dendra2 into various cell lines and populations. For our H1299 NSCLC (with pal-Dendra2) and myeloma (using H2B-Dendra2) lines, the cells were stably transduced with their respective Dendra2 plasmids using standard 2^nd^ generation lentiviral transduction methodologies; more extensive protocol information can be found online at Addgene and the Trono lab websites^50,51^. For our 4T1 mouse mammary carcinoma line, we introduced H2B-Dendra2 via a non-viral DNA transposon system (Sleeping Beauty) for stable integration into the genome; for cell populations and subpopulations resistant to viral transduction, a transposon system allows for the stable expression using any non-viral transfection delivery method^52,53^. After stable introduction of Dendra2, sort Dendra2 positive cells with homogeneous Dendra2 expression using FACS. Distinct cell lines can have different transfection efficiencies and the lines used ranged from 20-80% (data not shown). This limitation could lead to a loss of heterogeneity or a shift in subpopulation percentages. Therefore, it is important to confirm the heterogeneity characteristics of the cell line of interest after PCFP introduction. For our SaGA experiments, we confirmed no significant difference in invasive properties, cellular circularity and morphology, subpopulation percentages and cell phenotype (data not shown). Regardless, less obvious cellular functions may not be preserved with the addition of the exogenous tag. Further information and additional techniques for PCFP introduction can be found in the literature^47,54,55^.

#### Tissue culture conditions

Three common culture conditions are often utilized to assess cell phenotype *in vitro* – non-adherent, 2D and 3D culture. Depending on the specific cell type and/or experimental question, SaGA can and has been successfully used in all three settings (Fig. 4A). SaGA can be implemented under non-adherent culture to determine a variety of biological phenomena including differential cell responses to a treatment (i.e., cytokine, growth factor, drug, starve/stimulation) or in heterotypic cell mixing experiments. For example, glioma cells lose their ability to grow diffusely in the brain when grown as adherent cells and, therefore, passage and analyses in suspension under non-adherent conditions provide an *in vitro* environment conducive to modeling that particular *in situ* behavior^56^. Traditional 2D monolayer (including growth on standard tissue culture plastic dishes) is a common approach to exploring a range of cell biology questions and provides a simpler environment to implement SaGA. Additionally, SaGA can be combined with 3D culture techniques with physiologically and pathologically relevant extracellular matrices (ECMs). 3D culturing techniques allow for active matrix and structural cell remodeling through a “dynamic reciprocity” between cell and environment that has been shown to more faithfully recapitulate tissue specific function compared to 2D conditions^57,58^. In our laboratory, SaGA has been utilized to observe invasive phenotypes in a variety of ECMs, with different cell lines, each having their own distinct invasion phenotypes (Box 1, Fig. 2A, F).

**Fig. 4.**
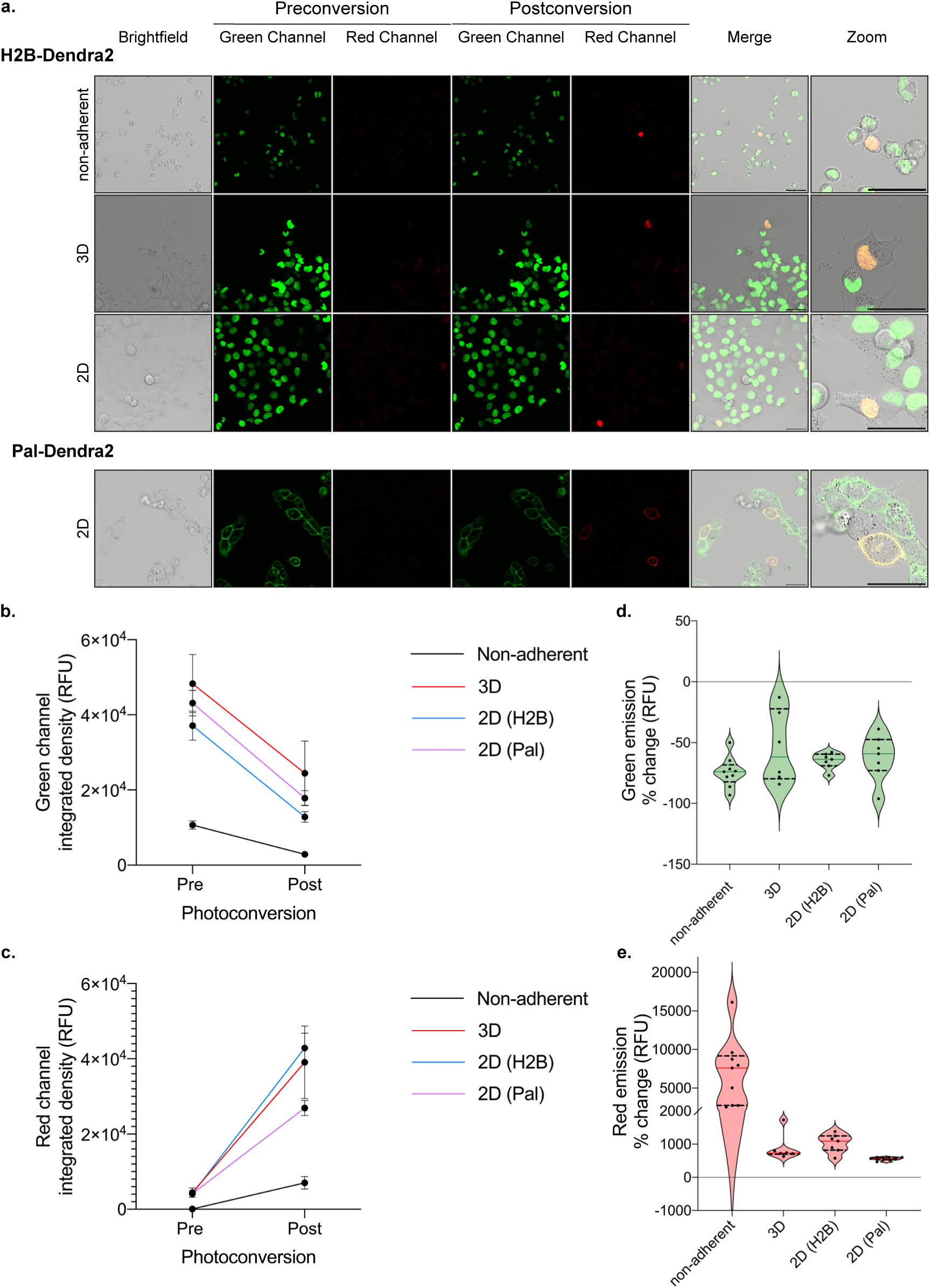
I Example photoconversion in different cell culture conditions. **a.** Cells stably expressing a photocon-vertible tag (ex: H2B-Dendra2, Pal-Dendra2) can be prepared under non-adherent, 3D, or 2D experimental conditions which illicit distinct and imageable cellular response for photoconversion. Non-adherent conditions were performed with RPMI8226 myeloma cells; H1299 lung cancer cells were used for all other conditions. Scale bar, 50 μm. **b, c.** Integrated density (relative fluorescence units) quantification of 6 or more cells pre-and post-photoconversion in the green (b) and red (c) channels, emission peaks, 507 nm, and 573 nm, respectively. **d, e.** Quantification of integrated density percent change of 6 or more cells pre-and post-photoconversion in the green (d) and red (e) channels.

##### Box 1

###### I 3D spheroid invasive area and circularity quantification

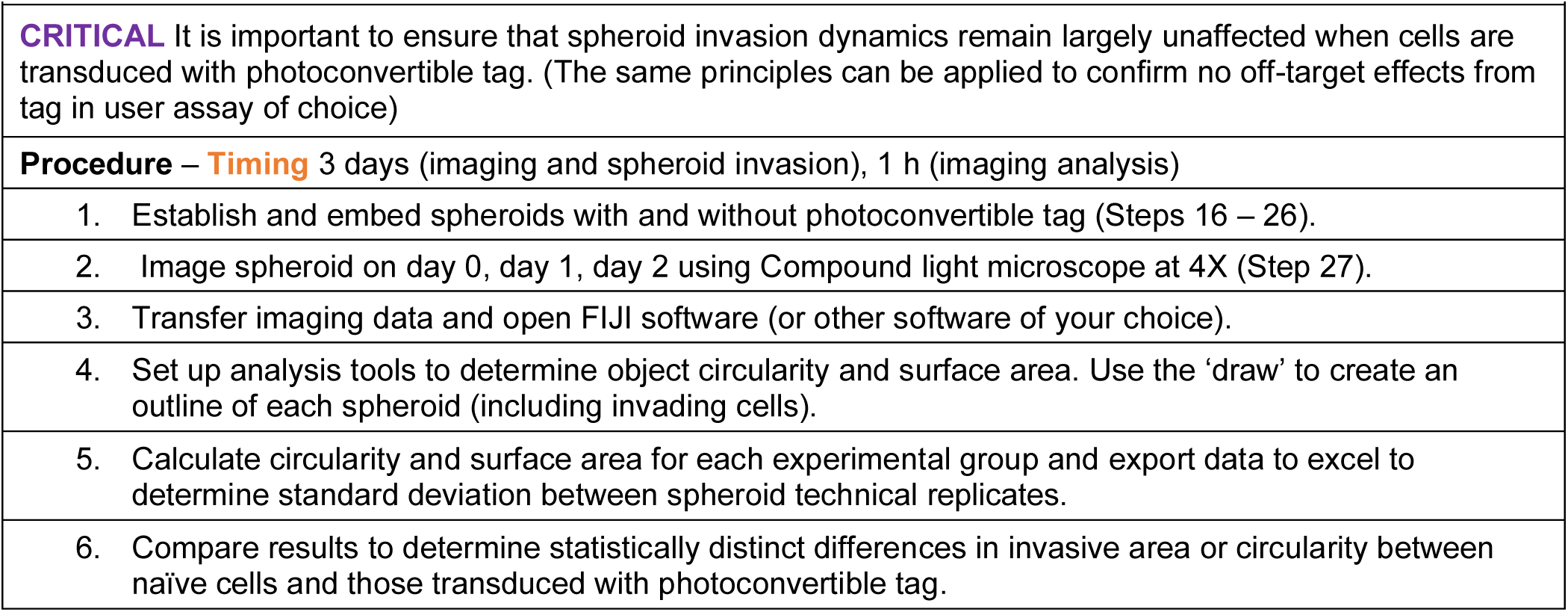

#### Live cell imaging and photoconversion

We conducted live cell imaging on a Leica TCS SP8 inverted point scanning confocal microscope equipped with a stage top incubator to maintain cell culture conditions while imaging. This microscope provides flexibility to modify laser intensity settings and includes a module for fluorescence recovery after photobleaching (FRAP, used for photoconversion here), where short laser pulses in a spatially defined region produce high energy light to induce spectral changes. FRAP has a longstanding history of being utilized to dissect protein dynamics and molecular diffusion, and many modern microscopes are programmed to include a pre-existing FRAP option^59–61^. Photoconversion uses a similar microscopy setup as FRAP, with the addition of a second (post-conversion) emission detection step; as such, the FRAP module may often be adaptable to the photoconversion step within SaGA. Within this module, a user can define a precise region of interest (ROI) that can vary from sub-cellular to multi-cellular in area (Fig. 4A). Microscopes with similar point scanning confocal, live cell capabilities, and appropriate laser lines, along with scanning-photoconversion modes, can be used to perform SaGA.

Dendra2 utilizes three laser lines for excitation and photoconversion: 405 nm or 488 nm (implement photoconversion), 488 nm (excitation pre-photoconversion, detection 490 – 550 nm), and 543-, 561-, or 568 nm (excitation post-photoconversion, detection 570 – 670 nm). Importantly, any additional small molecule or antibody dual-labeling is limited to the far-red spectrum^45^. Dendra2 photoconversion results in an approximate 50% decrease in the green channel and greater than 500% increase in the red channel, independent of culture conditions (Fig. 4B-E). Further, photoconversion of Dendra2 (Dendra2-green) is irreversible and after 14 hours we still observe photoactivated Dendra2 red fluorescent signal (Dendra2-red) utilizing our current optical settings. If the experimental procedure requires more than 24 hours between photoconversion and fluorescence activated cell sorting, the PS-CFP2 PCFP has been shown to remain stable for 48 hours and can be used as an alternative option^62^.

Insufficient photoconversion due to inadequate excitation light and imaging parameters can yield poor photoconversion efficiency. Notably, low photoconversion efficiency can lead to poor separation between non-photoconverted cells and photoconverted cells during FACS, which can compromise sorted cell purity. Conversely, photobleaching or phototoxicity may result from excessive laser power or excitation time. Photoconversion using illumination with the 405 nm laser line may be best resolved in short pulses with low laser intensity to avoid DNA damage (as UV damage disrupts nuclei division^63^). Alternatively, the 488 nm laser can be applied for more continuous pulses at low or moderate intensity; for this modification, the decreased efficiency of the photoconversion stimulated by a 488 nm laser (compared to the 405 nm laser) requires an increase in excitation duration^45^. Phototoxicity from high intensity laser exposure is a limiting experimental factor^64^. Dead cells can be identified and avoided by staining with a live/dead stain during FACS sorting (Fig. 5A). Similarly, since Dendra2 can be photoconverted while simultaneously visualizing the green pre-converted fluorophore at 488 nm, it is important that the laser power is reduced when visualizing the Dendra2-green for ROI identification to avoid unwanted photoconversion.

**Fig. 5.**
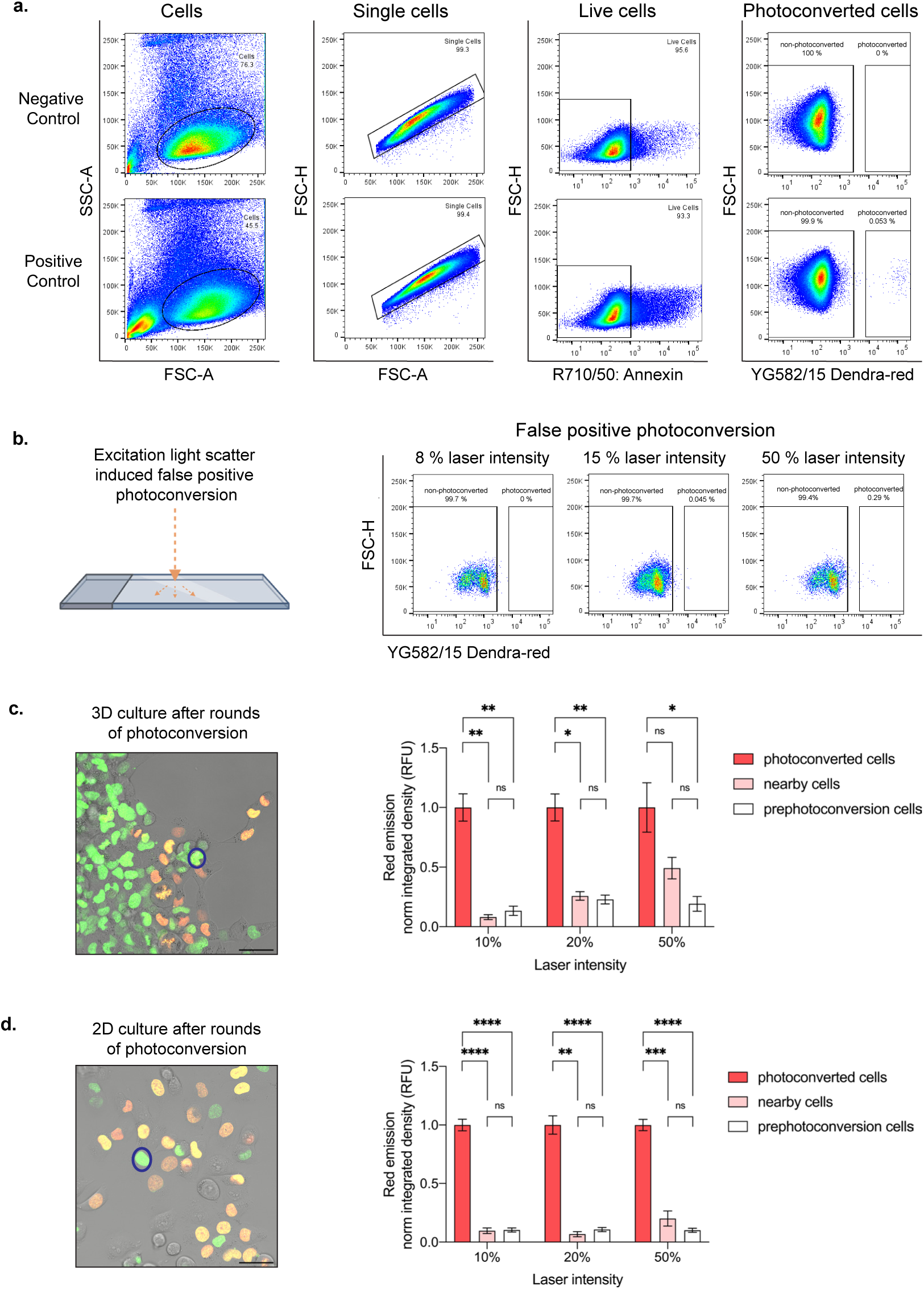
I Cell selection and isolation optimization. **a.** Flow plots illustrating stepwise isolation of live photocon-verted cells. 8 % 405 nm laser line intensity utilized in positive control. **b.** False positive photoconverted cells due to light reflection off the glass plate at varying photoconversion laser intensities at 405 nm. **c.** Representative merged image showing photoconversion of multiple cells (orange and yellow cells) in 3D, where intensity change is measured in a neighboring, non-photoconverted cell (representative nearby cell,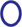). Quantification showing fold change of normalized red emission after rounds of photoconversion are complete. **d.** Representative merged image showing photoconversion in multiple cells (orange and yellow cells) in 2D, where intensity change is measured in a neighboring, non-photoconverted cell (representative nearby cell,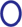). Quantification showing fold change of normalized red emission after rounds of photoconversion are complete. *p < 0.05 by one-way ANOVA with Tukey’s multiple comparisons test. Scale bar, 50 μm.

Additionally, optimizing for a conservatively defined ROI reduces off-target and false positive nearby cell photoconversion. To determine the potential of false positive and off-target photoconversion to occur, we performed two troubleshooting experiments (Fig. 5B – D). First, to test whether the reflection of light has the capacity to photoconvert adjacent cells, we applied the 405 nm laser line at varying intensities to an empty ROI surrounded by cells and then performed FACS to determine percent of photoconverted cells (Fig. 5B). These experiments resulted in false positive photoconversion only when 50% laser line intensity was used, well above the range of photoconversion intensity values used in our system (5 – 15 %) (Fig. 5B, Box 2). Next, we tested the impact of multiple rounds of photoconversion on nearby, non-photoconverted cells within a single field-of-view by measuring their Dendra2-red emission after photoconverting 10 or greater adjacent cells (Fig. 5C, D). This experiment was conducted in 2D and 3D culture conditions utilizing varying degrees of laser line intensities (Fig. 5C, D). After rounds of photoconversion under 3D culture conditions within a single field of view, we visualized no significant change in the nearby cells’ red emission with 10% 405 nm laser intensity conditions and little to no change with 20% laser intensity (Fig. 5C). However, when utilizing the 405 nm laser at 50% intensity, the Dendra2-red fluorescent signal of nearby cells increased drastically and resulted in no significant difference when compared to the red fluorescent signal of photoconverted adjacent cells (Fig. 5C). These data suggest that photoconversion at 50% 405 nm laser line intensity can lead to the collection of falsely photoconverted cells, unlike cells photoconverted at 10% or 20% laser line intensities. Interestingly, after rounds of photoconversion under 2D conditions, we observed no significant change in the nearby cells’ Dendra2-red emission when utilizing 10%, 20%, or 50% laser line intensities (Fig. 5D). It is important to note that the percent laser intensity will vary by microscope and/or system conditions, therefore, intensity values should be determined independently. Overall, the optimal approach is to utilize the lowest laser intensity and exposure time that can successfully photoconvert cells for efficient live cell sorting.

##### Box 2

###### I Photoconversion time course guidelines

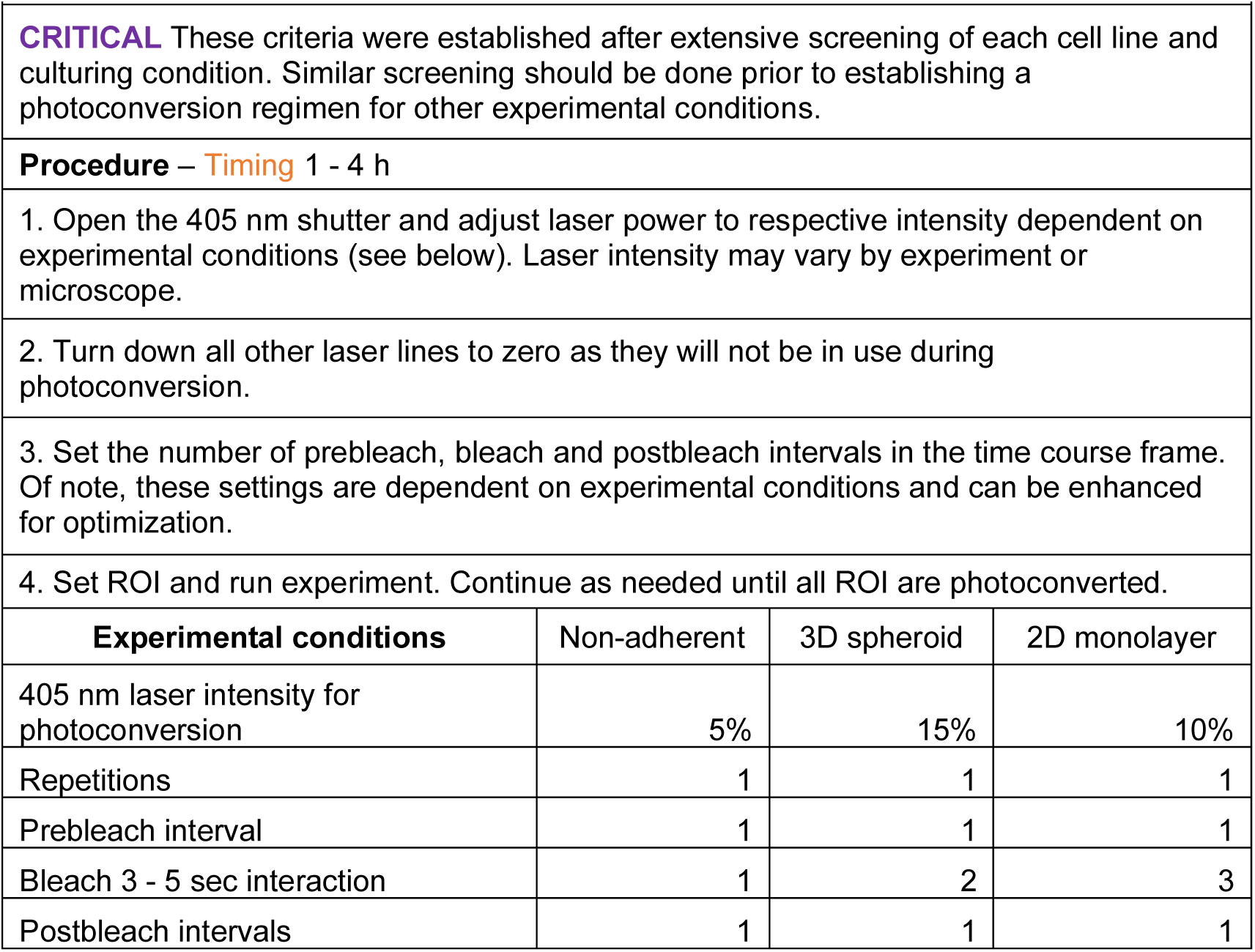

#### Fluorescence activated cell sorting

A fluorescence activated cell sorter (FACS) with a minimum requirement of two-color flow cytometry is used to isolate user-defined cells based upon fluorescent state. With Dendra2, in each sample, the green cells include one or more phenotype(s), and the red cells include the photoconverted single or set of cells photoconverted based on phenotype. We typically ensure at least 50 cells or greater are photoconverted to account for cell loss due to viability or, in the case of 3D samples, matrix degradation and cell retrieval steps. For 3D collective invasion experiments, multiple follower and leader cells were separately photoconverted, harvested from their ECM and sorted for downstream applications. In all experiments, negative controls were used to set initial gating: Dendra2 positive cells not exposed to 405-or 488 nm laser lines to determine autofluorescent signal emission in the red channel (yellow green laser line (YG), bandpass filter: 582/15 Dendra2-red), live/dead cell staining to collect only live cells and avoid dead cell autofluorescence, and Dendra2 cells exposed to 405 nm laser line, but purposely not photoconverted to determine percent of cells falsely photoconverted (these cells were exposed to the same photoconversion time course as the positive control) (Fig. 5A). Non-photoconverted cells were detected for emission within the Dendra2-red channel to determine the rate of false positives, and photoconverted cells were detected and gated for sorting within the red channel (Fig. 5A). FACS gating for the collection of photoconverted cells relied on the detection of separate events (Fig. 5A). Sorted populations are typically 95% or more in fluorescent purity; however, samples may include false positives. To ensure FACS purity, a 100-fold separation between non-photoconverted (based upon gating parameters set by the negative control) and photoconverted (Dendra2-red positive) cells is desired (Fig. 5A). Photoconversion parameters can be altered to optimize sorting efficiency. Similarly, we recommend performing a trial experiment where a defined number of cells are photoconverted and sorted to test SaGA platform efficiency. After sorting, cells for further phenotypic analysis were replated into complete growth medium for long-term cellular cultivation and propagation. Cells for immediate multi-omic profiling were pelleted, flash frozen, and then stored in a negative 80 °C freezer.

After utilizing FACS to isolate Dendra2-red photoconverted cells, there are a variety of possibilities for downstream analysis to determine the molecular significance of the phenotypically isolated cell subpopulation. For example, to determine whether the cells of interest contain additional heterogeneity, photoconverted single cells can immediately be dropped into a 96-well plate for single cell -omic analysis (Fig. 6, Table 1). Cell subpopulations can also be submitted for bulk multi-omic analysis like RNA sequencing or DNA methylation array (Table 1). Likewise, to determine whether the isolated population(s) maintains its respective phenotype over a series of passages (i.e., phenotypic stability), cells can be cultured for further downstream analysis. Together, these downstream analyses provide multi-dimensional molecular depth to the phenotypic distinctions.

**Fig. 6.**
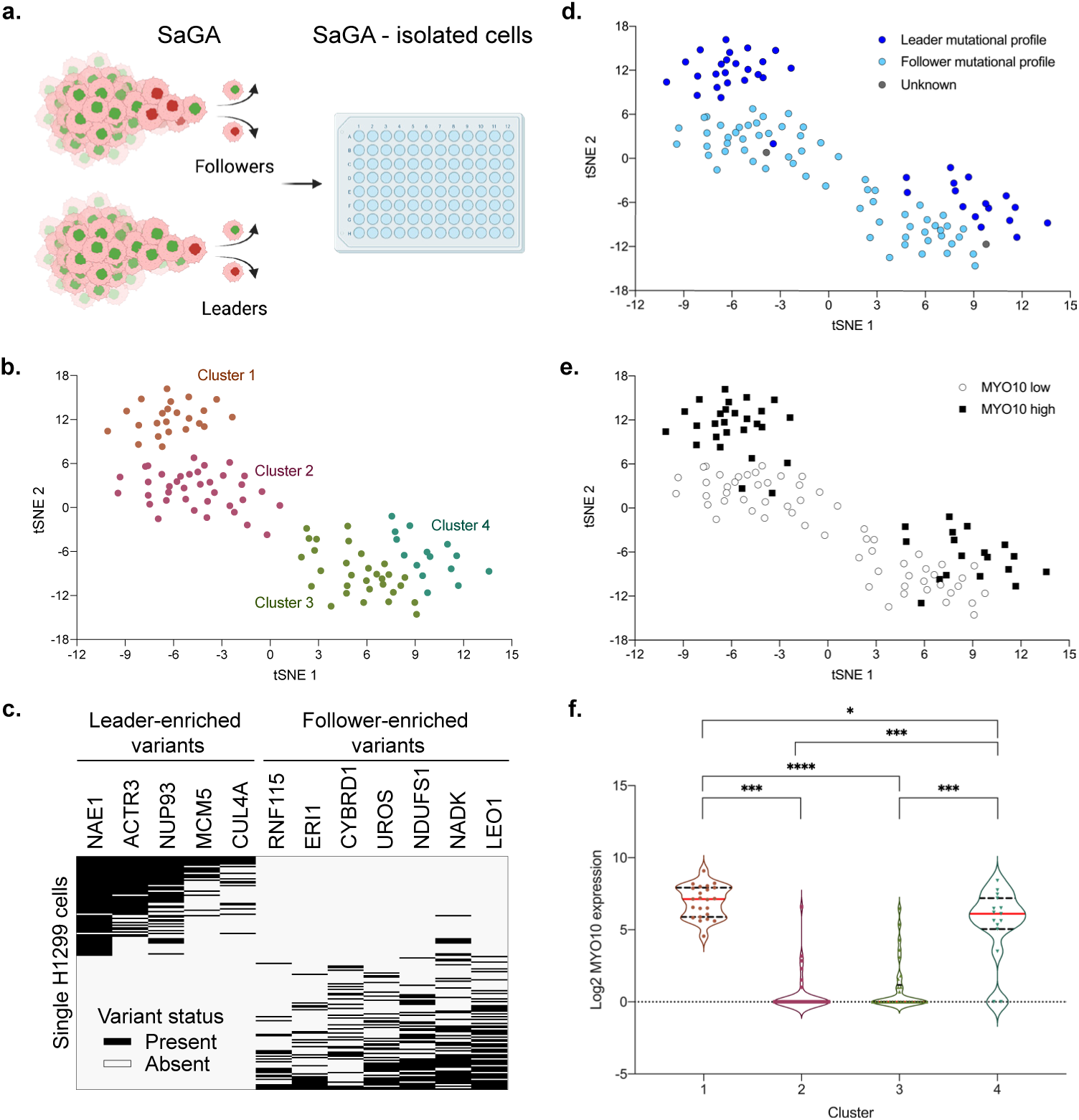
I SaGA-isolated single cell RNA sequencing and analysis. **a.** Workflow schematic of collection of SaGA-isolated cells for single cell RNA sequencing and analysis. H1299 cells were used to perform a 3D spher-oid invasion assay for a 24 h invasion period. Upon application of SaGA, user-defined cell populations (Leaders and Followers) were sorted directly into a 96-well plate with RNA lysis buffer, frozen at – 80 °C, and subsequently processed for single-cell RNA sequencing. Created with Biorender.com. **b.** tSNE plot of single H1299 cells (n = 105) based upon expression of the most variably expressed genes (n = 1,155). tSNE clusters were determined by cell positioning within the plot. **c.** Mutation profile plot for n = 171 single cells using previously identified H1299 leader-and follower-specific mutations^30^. **d.** tSNE plot with each cell labeled by its mutation profile from panel c. **e.** tSNE plot with each cell labeled by high (defined as [log2 (normalized counts + 1)] >2 or low <2 expression of Myo10. **f.** Quantification of Myo10 expression for each tSNE cluster. *p < 0.05 by one-way ANOVA with Tukey’s multiple comparisons test.

### COMPARISON WITH OTHER METHODS

Cell subpopulation heterogeneity can be assessed using several approaches, including flow cytometry analysis, live-cell imaging, and single-cell multi-omics. However, these methods have limitations since they are unable to directly link live cellular phenotypes, geographic information, and molecular signatures. The SaGA platform leverages multiple experimental modalities to generate multi-scale datasets that integrates molecular, phenotypic, and spatial data.

FACS is a technique initially developed for immune cell classification, and now is utilized in a variety of fields to identify subpopulation heterogeneity within a larger population^65–67^. Traditional approaches utilizing FACS often isolate subpopulations with known markers, making it difficult to identify novel subpopulations utilizing FACS alone. SaGA takes advantage of the ability of FACS to isolate fluorescent single cells and couples that with the preservation of historical spatial and phenotypic information. Importantly, SaGA combines live-cell imaging with FACS to enable propagation of the isolated cells, which we have shown can maintain stable phenotypes over time^26^.

These spatially defined and isolated cells can be further analyzed by multi-omics to evaluate molecular signatures and define novel subpopulations. For example, single-cell RNA sequencing has led to the resolution of small, rare subpopulations within the bulk population^68^. However, the power of this approach is limited by the collection process of the cells. Spatial localization is lost upon dissociation of cells for single-cell sequencing. Consequently, single-cell sequencing data alone does not provide insight to phenotypic or spatiotemporal distinctions within the established subpopulations. Since SaGA allows for the precise isolation of cells based on phenotype of interest, it adds context to the datasets driven by multi-omic platforms. By using a photoconvertible approach for cell selection, SaGA can be applied to many different contexts, depending on the interest of the researcher. This is an advantage over selecting cells using microfluidic systems. While the field of microfluidics has evolved to incorporate several methods to distinguish cells and isolate cell subpopulations^69,70^, these techniques are still limited in their capacity to maintain spatial information, as cells are applied to these systems in suspension. It would be difficult to isolate adherent cells of interest or cells invading from a spheroid using a standard microfluidic, in contrast to SaGA^26^. Additionally, SaGA can be performed using equipment commonly available to researchers in the biomedical field without requiring expertise to engineer the microfluidic.

The recent advancements of spatial multi-omic methodologies allow for comprehensive assessment of molecular phenotypes in tissue while retaining spatial tissue context^17,71–77^. For example, spatial transcriptomics can assess global gene expression patterns and integrate these data with positional localization of cells^75,78,79^. While these methods identify different cell populations within a single tissue and maintain the tissue’s architecture, the samples are frozen or fixed. SaGA is advantageous in that the samples are viable throughout the entire process and a historical live cell phenotype can be integrated. Cells are identified and isolated during live-cell microscopy and the purified population can be grown for long-term cultivation with traditional tissue culture techniques. This allows for either immediate sequencing analysis or analysis after long-term propagation. The populations isolated by SaGA can be analyzed by a myriad of live-cell assays depending on the user’s interest, not limited by experimental conditions (i.e., fixed tissue or in suspension).

### OVERVIEW OF THE PROCEDURE

Here, we exploit phenotypic variations within a parental cell population to isolate and characterize distinct cellular subpopulations. Our protocol begins with the preparation of cells in non-adherent, 2D, and 3D *in vitro* environments (Steps 1 – 29). Next, we select and isolate phenotypically distinct cell(s) utilizing live confocal microscopy and FACS (Steps 30 – 69). Lastly, we provide multi-omic or phenotypic modes of analyses for the SaGA-isolated subpopulations to determine unique biological phenomena (Steps 70-72).

## MATERIALS

### Reagents

Biological

— H1299 and RPMI8226 cell lines purchased from American Type Culture Collection, cat. nos. CRL5803 and CCL-155, respectively.
— Lenti-X 293T cells purchased from Clontech Labs cat no. 632180. **! CAUTION** Authenticity of cell lines should be validated via STR profiling. Sterility of cell lines should be maintained by appropriate and routine checks; these include morphological checks by microscopy, and mycoplasma contamination checks by Myco Alert Mycoplasma Detection Kit (or similar).

Technical

— Accudrop Beads (BD Biosciences cat. no. 345249)
— Annexin V, Pacific Blue Conjugate (Thermo Scientific cat. no. A35122)
— Annexin V, Alexa Fluor 680 (Thermo Scientific cat. no. A35109)
— Collagen I, High Concentration, Rat Tail, 100 mg (Corning cat. no. 354249)
— Collagenase/Dispase Roche (Millipore Sigma cat. no. 11097113001)
— CS&T Research Beads (BD Biosciences cat. no. 650621)
— DPBS, no calcium, no magnesium (Gibco cat. no. 14190250)
— Fetal Bovine Serum, Regular, USDA Approved (Corning cat. no. 35-011-CV)
— Fetal Bovine Serum, Dialyzed, US Origin (Thermo Fisher Scientific cat. no. 26400044)
— Growth Factor Reduced Basement Membrane Matrix (Corning cat. no. 356231)
— Lipofectamine 3000 Transfection Reagent (Invitrogen cat. no. L3000001)
— MycoAlert Mycoplasma Detection Kit (Lonza cat. no. LT07-118)
— OptiMEM I (Gibco cat. no. 31985062)
— Palmitoylated Dendra2, (Emory University, Gary Bassell Lab)
— Penicillin Streptomycin (10,000 U/mL) (Thermo Fisher Scientific cat. no. 15-140-122)
— pLenti.CAG.H2B-Dendra2.W Plasmid (Addgene cat. no. 51005)
— pMD2.G (Addgene cat. no. 12259)
— Polybrene (Millipore cat. no. TR 1003-G)
— psPAX2 (Addgene cat. no. 12260)
— RPMI 1640 [-] L-glutamine, Phenol Red (Gibco cat. no. 11875101)
— RPMI 1640 [-] L-glutamine Medium, no Phenol Red (Gibco cat. no. 11835030)
— Sterilized Distilled Water
— Trypan Blue Solution 0.4% (Sigma-Aldrich cat. no. T8154)
— Trypsin/EDTA Solution (Lonza cat. no. CC-5012)
— Trypsin Neutralizing Solution (Lonza cat. no. CC-5002)

### Reagent Set Up

— Complete medium – Prepare RPMI 1640 (with and without phenol red) supplemented with 10 % (vol/vol) FBS and 1 % (vol/vol) Penicillin Streptomycin. Prepared media can be stored at 4 °C for two weeks. Maintaining sterile culture, remove necessary volume as needed and pre-warm aliquot to 37 °C before use (unless otherwise noted).
— Flow buffer – Prepare fresh on the day of use by supplementing RPMI 1640 or appropriate base media (without phenol red) with 5 % (vol/vol) dialyzed FBS. Alternatively, calcium free reagents can be used to reduce cell aggregation.

#### 3D matrix embedment experiments

**CRITICAL** Particularly critical for epithelial and endothelial cell differentiation and function, the reconstituted basement membrane (rBM) provides a critical structural interface between certain cell types and their surrounding environment, acting as a mechanical buffer and barrier to both cellular and molecular traffic^80,81^. rBM derived from Englebreth-Holm-Swarm murine tumors provides a laminin and type IV collagen rich microenvironment conducive to studying 3D cell growth and differentiation, as well as morphogenesis (breast, lung) and invasion. As such, rBM provides a microenvironment conducive to collective invasion for many of our lung tumor cell lines (Fig. 2) ^82,83^. Conversely, for our studies using the 4T1 murine model of breast cancer, we find that fibrillar type I collagen provides a microenvironment supportive of multicellular pack invasion for the largely e-cadherin positive 4T1 cells (data not shown).

— Recombinant basement membrane master mix – Thaw growth factor reduced recombinant basement membrane (often referred to as its tradename Matrigel, rBM) at 4 °C overnight and maintain on ice to avoid polymerization at room temperature. Spheroids are embedded in 5 mg/mL rBM (in full media) and then plated onto a glass bottom dish. Note, the polymerization characteristics of rBM may change from lot to lot of Matrigel; as such, the concentration may need to be adjusted on a lot-specific basis. Additionally, cell secreted growth factors and cytokines will vary between lots of non-growth factor depleted Matrigel; growth factor reduced Matrigel leads to a more consistent composition^84^.
— Collagen I master mix – Keep stock collagen I (high concentration derived from rat tail) solution at 4°C or on ice until ready for use. On ice, supplement stock collagen I with 10 % (vol/vol) PBS to a working concentration of 3 mg/mL. Check pH of solution with pH strips and adjust pH to approximately 7.0. Keeping working solution on ice until ready to use. Do not store dilute solution long-term.
— Collagenase/dispase cocktail: Suspend 100 mg of collagenase/dispase (C/D) in 1 mL of PBS or base media (without added growth factors or FBS) for a 100 mg/mL stock solution. Importantly, calcium is an important factor for enzymatic stability and activity in this cocktail; therefore, in the case of PBS, calcium supplemented PBS is required. Pipette gently to thoroughly resuspend. Sterilize with a 0.2 μm filter. Make aliquots of stock at 50-100 μL per aliquot. Freeze at ࢤ20 °C until ready for use.

### Equipment

Tissue culture

— CellDrop Automated Cell Counter (DeNovix cat. no. CellDrop FL-UMLTD)
— CO_2_ Incubator Forma Series II Water Jacketed (Thermo Fisher Scientific cat. no. 3110)
— Cryostorage Container, Locator 4 Plus (Thermo Fisher Scientific cat. no. CY509108)
— Eppendorf Centrifuge 5425 (Thermo Fisher Scientific cat. no. 13864455)
— Eppendorf Centrifuge 5810/5810R (Millipore Sigma cat. no. EP022628168)
— Falcon Round-Bottom Tubes with Cell Strainer Cap, 5 mL (Stem Cell Technologies cat. no. 38030)
— Isotemp Dual Digital Water Bath (Fisher Scientific cat. no. FS-215)
— Sterilized Biosafety Cabinet (Labconco cat. no. 3440009)
— Ultra-Clear Microcentrifuge Tube 1.7 mL (DOT Scientific cat. no. 609-GMT)
— Ultra-Low Attachment Multiwell Plates, Sterile (Corning cat. no. 29443-034)
— µ-Slide 8-Well Glass Bottom (Ibidi cat. no. 80827)
— 15 mL Conical Centrifuge Tube (VWR, cat. no. 430052)
— 35 mm Glass Bottom Dish (MatTek, part no. P35G-1.5-14-C)
— 50 mL Conical Tube (VWR cat. no. 430290)
— 75 cm^2^ U-shape cell culture flask, canted neck (Corning cat. no. CLS430641U)

Imaging and FACS

— BD FACS Aria II Cell Sorter
— Inverted Microscope, Olympus CKX41
— Leica TCS SP8 Inverted Point Scanning Confocal equipped with Galvano and 8 kHz resonant scanners, Tokai Hit stage top incubator for CO2 and temperature control, two multi-alkali PMTs, two HyDs, and one transmitted light PMT

Software

— BD FACSDiva Software
— FlowJo
— Graphpad Prism
— ImageJ or FIJI
— LAS X Imaging Software

## PROCEDURE

### Sample Preparation

**CRITICAL** This section describes our approach to handling a variety of cell lines and patient samples in culture. Non-adherent and 2D conditions are respective to those particular and potentially uniquely appropriate to the experimental questions and cell treatments. 3D tumor spheroid formation is described in detail by using the example human NSCLC H1299 cell line. See Table 2 for parameters to consider for optimizing the 3D invasion assay procedure to other cell lines and systems.

**Table 2.** I Parameters for spheroid formation

| Table 2 \| Parameters for spheroid formation |  |  |  |
| --- | --- | --- | --- |
| Parameters | Optimization | Examples | Steps |
| Seeding density | Spheroid density depends upon cell size, shape, morphology, and rate of proliferation. Various densities should be screened to achieve ~ 500 µm in spheroid diameter upon embedding into matrix. Importantly, the larger the cell number, the larger the spheroid, the greater oxygen differential between the external cells and the cells internal to the 3D structures. | 3000 cells/well (H1299, 4T1)<br>1000 cells/well (A375) | Steps 16 – 18 |
| Nanoparticle contamination | Sterile 96-well plates and/or sterile pipet tips are often contaminated with sterilized nanoparticles that can become embedded within a spheroid and deform its shape. 1.5X spheroids are created to account for unusable spheroids. |  | Steps 16 – 25 |
| Cell adherence | Heterogeneous cells can express distinct adherence junction profiles to regulate cell – cell junctions and cell – matrix adhesion/interactions. Centrifugating 96-well plate places cells in the center of the well near one another to promote cell – cell junction formation. Upon spheroid formation, different matrices can be screened to determine the ability for cell – matrix adhesion formation and interactions. | rBM (H1299, A375)<br>Collagen I (4T1) | Step 18 |
| Time | After centrifugation, spheroid formation requires a 24 h or more incubation time. Cells can be screened to determine optimal incubation time to maintain both spheroid integrity during the embedding process and cell viability after. | 72 h (H1299, 4T1, A375) | Step 18 |

### Thawing and maintenance of cells

**Timing** 1 h (thawing cells), 1 week (cell growth and maintenance)

1. Prepare cell culture media as described in reagent set up.
2. Warm desired volume of culture media in 50 mL conical tubes at 37 °C in bead or water bath.
3. Prepare tissue culture appropriate laminar flow hood using UV light and wipe down all working surfaces with 70 % ethanol (EtOH, vol/vol). Perform all cell culture within the laminar flow hood to maintain sterility. Use rigorous aseptic/sterile tissue culture technique where appropriate.
4. Rapidly thaw a vial of RPMI8226-Dendra2 or H1299-Dendra2 (either pal-or H2B) cells by gentle agitation in 37 °C water bath.
5. Decontaminate the vial by spraying with 70 % (vol/vol) EtOH.
6. In the tissue culture hood, quickly open and transfer the vial contents to a 15 mL centrifuge tube containing 9.0 mL complete culture medium and spin at 125 g at room temperature (RT) for 5 min to pellet the cells out of solution. Speed and times may vary between cell lines.
7. Carefully remove the supernatant from above the pellet, taking care not to disturb the pellet. Resuspend the cell pellet with complete media and dispense into a 25 cm^2^ or a 75 cm^2^ (if working with cell concentrations greater than 1 x 10^6^) culture flask. The density of the cells and volume of culture media may vary between cell lines.
8. Incubate at 37 °C and atmosphere of 95 % Air, 5 % CO_2_. These conditions may vary by cell line.
9. Optional: when necessary, change media after 24 h to eliminate cells that do not survive thaw cycle.
10. Allow cells to recover from thaw (typically 2-3 days) and passage 1X prior to experiment set up.
11. Ensure media renewal occurs once every two days, or as appropriate for the cell line.
12. Passage cells at less than 70 % confluency and ensure to collect all cells. **CRITICAL STEP** With heterogenous cell lines and patient samples, under normally adherent conditions, floating cells may be viable, non-adherent subpopulations within the greater population. When necessary, collect the floating cells, as well as the difficult to de-adhere cells, to ensure that the overall population doesn’t artificially drift in subpopulation composition.

a. For adherent cell passaging:

i. Collect conditioned/spent media in 50 mL conical tube.
ii. Wash 1X with PBS and add to the conditioned media.
iii. Detach cells with method of choice (trypsin, EDTA, Accutase). Different cell types may require distinct cell detachment mechanisms; therefore, it is best to determine the best passaging conditions for your system.
iv. Neutralize trypsin enzyme with an appropriate trypsin neutralizing solution (TNS) at a 1:1 trypsin to TNS ratio. While some TNS product utilize serum (FBS or otherwise) to provide excess substrate for the trypsin to enzymatically digest protein molecules, an alternative – particularly for cells cultured serum free – is the use of a TNS product consisting of a 1X PBS solution containing 0.0125 % (vol/vol) soybean trypsin inhibitor.
v. Pipet additional complete media onto plate to wash and collect all cells from plate and add to the 50 mL conical tube.
vi. Centrifuge conical tube at 125-250 g RT for 5 min, centrifuge speed depends on the cell line.
vii. Resuspend in 3 mL of complete media.
viii. Count cells using 0.4 % Trypan blue solution at a 1:1 ratio with hemocytometer.
b. Non-adherent cells do not require enzymatic cleavage from cell plate and can simply be collected from plate at time of experimentation. Cell concentrations may vary by plate or cell size.

### Option A: Non-adherent sample preparation for photoconversion

**Timing** 1 h (cell collection, counting and plating)

13. Collect and count RPMI8226-Dendra2 cells using 0.4 % Trypan blue at a 1:1 ratio with hemocytometer.
14. 0.5 × 10^6^ cells are isolated from the bulk sample and spun down at 125 – 250 g RT for 5 min, centrifuge speeds may vary by cell line. Number of cells will vary by plate and cell size.
15. Resuspend cells in 200 µL phenol-red free complete media and transfer to 8-well glass bottom slide chamber. Cell number is dependent on cell morphology and well dimensions. **CRITICAL STEP** Cells in suspension will gravitate down to the bottom of the well over time. To avoid cell plane variability and maintain a relatively constant z – plane when imaging, incubate the cells at least 30 min in a static environment prior to photoconversion. These cells will continue to move across the X, Y and Z planes over time and microscope parameters should be adjusted accordingly.

### Option B: 3D cell culture sample preparation for photoconversion

**Timing** 1 h (cell plating), 72 h (incubation), 2 h (embedding into matrix), 24 – 48 h (incubation for invasion overtime)

**CRITICAL** During processes such as morphogenesis and tumor progression, a 3D microenvironment provides a physical and biochemical scaffold for cells to grow, self-organize and remodel surrounding pericellular and extracellular matrices. *In vivo* and *in situ*, 3D matrix microenvironments are comprised of cellular (fibroblasts, lymphocytes, endothelial cells, etc.) and non-cellular (ECM, cytokines, growth factors) components that actively mediate cell migration and invasion, proliferative capacity, and differentiation^85–87^. Distinct ECMs are highly specialized and organized networks integrating a myriad of molecules that establish an essential architectural support structure for cells, tissues, and organs along with providing a substrate for cell adhesion and traction ^81,88^. Various 3D *in vitro* systems utilize reconstituted ECMs to assess growth, motility, and invasive phenotypes. Depending on cell type, biological activity of choice (such as single cell and collective invasion), different ECMs may be better suited to asking specific biological questions.

16. To generate spheroids, plate 3,000 H1299 cells in 200 μL (1.5 × 10^4^ cells per mL) in a low adherence 96-well plate. Create 18 spheroids per condition to embed 10 – 12 spheroids in matrix. Excess spheroids are created to account for shape and stability deformities during incubation period for spheroid formation (Table 2). The cell number may be adjusted for different cell lines and spheroids of different diameters (Table 2).
17. Centrifuge plate at 450 g RT for 10 min to collect cells at the center of each low absorbance round-bottom well.
18. Incubate cells for 72 h for cell – cell junction formation within the spheroid. **CRITICAL STEP** Some cells may resist adhering to one another to form spheroid. Incubation time varies on the cell line and in some instances, during the 72 h spheroid formation period, a once a day, 10 min at 450 g RT spin may encourage spheroid formation (Table 2). **? TROUBLESHOOTING**
19. Prior to the day of embedment, snip an experimentally appropriate number of 1,000 µL and 200 µL pipet tips for embedding spheroids into matrix. Autoclave to re-sterilize tips and allow to cool to room temperature.
20. A standard light microscope can be used prior to spheroid removal from the round bottom plate to validate spheroid integrity has been maintained (no debris contaminating or altering spheroid) (Table 2). Depending on the cell line, spheroids may not be spherical in shape. While sphericity is not necessary for transfer, spheroid stability is necessary for extraction from the round bottom plates and successful embedment. Non-stable spheroids/structures may fall apart during the process.
21. Collect usable spheroids with tip-snipped 1,000 µL pipet tips into 1.7 mL microcentrifuge tube - one experimental group per tube. The same snipped-tip can be used for all spheroids within the same experimental group. Pipet 8 spheroids per 1.7 mL microcentrifuge tube. The number of spheroids included can be increased or decreased depending on the experiment.
22. Allow the spheroids to sink to the bottom of the microcentrifuge tube. Remove excess media taking care not to pipet up the spheroids. Remove single and detached cells by washing spheroids with 1 mL of complete medium 2X by pipetting media along the edges of the tube in a circular motion, allowing the spheroids to sink, and removing excess media. **CRITICAL STEP** Treat spheroids with care. Avoid shaking tube, do not vortex. Spheroids are large and solid enough to visualize as they drop to the bottom of the microcentrifuge tube.
23. Collect spheroids in 100 µL 3D master mix using tip-snipped 200 µL pipetmen and plate in 35-mm glass bottom dish.
24. Use a second unsnipped 200 µL pipette tip to carefully spread master mix to cover entire glass surface.
25. With a third unsnipped 200 µL pipette tip, carefully move and spread spheroids for equal distribution. **CRITICAL STEP** If there is an air bubble in matrix, 70 % EtOH (vol/vol) can be used to eliminate the air bubble. Dip a clean 200 µL pipette tip into ethanol and poke the bubble. The alcohol breaks the surface tension of the matrix, causing the bubble to pop.
26. Allow >30 min for matrix to polymerize. After complete polymerization, add 1.5 mL pre-warmed complete medium and incubate. Note that the addition of cold media to the temperature-sensitive polymerized rBM may destabilize the matrix, leading to the loss of matrix and spheroids from the surface into the media.
27. Image every 24 h if monitoring surface area or circularity (Box 1).

### Option C: 2D cell culture sample preparation for photoconversion

**Timing** 1 h (cell collection, counting and plating)

28. Passage cells at < 70 % confluency and plate cells on glass bottom plate for photoconversion the following day. Cell number will be dependent on plate size and cell morphology. Cell confluency will be dependent on experimental question.
29. If distinct phenotypes are present temporally upon treatment with drug or other additive factor, adjust time course and confluency to match these criteria.

### Confocal imaging and photoconversion

**CRITICAL** This section describes the set-up and application of the Leica TCS SP8 inverted scanning confocal microscope for photoconversion during live cell imaging. We discuss a few Leica systematic anomalies; however, all steps can be adapted to fit the technical set up for most scanning confocal microscopes. The protocol by Chudakov, D.M., *et al*. provides specifics on Dendra2 photoconversion utilizing either the 405 nm or 488 nm laser lines and can be referenced for more detail^45^. The protocol we describe here uses the 405 nm laser line.

### Microscope and laser set-up

**Timing** 2 h (stage top incubator equilibration), 0.5 h (cells equilibrating to stage top incubator conditions)

30. Prepare stage top incubator to maintain standard tissue culture conditions, typically 5 % CO_2_ at 37 °C. These conditions may vary by cell line and are utilized to facilitate typical cellular phenotypes under defined experimental conditions.
31. Fill stage top incubator with enough autoclaved distilled water to maintain humidity and compensate for evaporation during imaging. Allow the incubator to equilibrate for at least 2 hours.
32. Turn on the computer, microscope, scanner, and laser power source. Photoconversion requires three laser lines: 405 nm for photoconversion, 488 nm for visualization of Dendra2-green (excitation and emission peaks: 490/507 nm), and 561 nm for visualization of Dendra2-red (excitation and emission peaks: 557/573 nm). Alternatively, can use 543-or 568 nm laser lines in place of 561 nm for visualization of Dendra2-red (emission spectral ranges: 570-670 nm).
33. Place the plate with cells inside stage top incubator and incubate for 30 min to allow cells to equilibrate to microscope conditions. **? TROUBLESHOOTING**
34. Open laser configuration window and turn on the Diode 405 nm (UV) laser, Argon laser (visible), and the DPSS 561 nm laser lines. **CRITICAL** Argon laser power intensity settings are specific to laser line and microscope conditions.
35. Image acquisition set-up.

(a) Turn off resonant scanning. Resonant scanning is a mode to increase imaging speed by gathering images at a rate of 30 frames per second or higher. This is typically used for overnight live cell imaging and therefore is not required for photoconversion.
(b) Set the scanning parameters to the XYT (XY Time) mode. Time is necessary for photoconversion (Step 37d). Z-plane is not required for photoconversion because high intensity laser exposure reaches multiple planes within the defined region of interest (ROI).
(c) Set scanning pixel size to be 1024 x 1024 or lower. Resolution can be sacrificed for increased imaging speed and pixel dwelling time within the ROI. High resolution is not required for successful photoconversion.
(d) Use the default scanning speed (400 – 800 Hz). This is the acquisition rate of pixels per second.
(e) Determine the zoom factor empirically. The zoom is dependent on the objective lens and user preference. A typical starting point is a zoom factor of 2 and adjust as needed based on ROI.
(f) Set line averaging at 2 and change as needed depending upon resolution needs or scanning speed requirements. **CRITICAL STEP** Scanning pixel size, speed, zoom factor, and line averaging are ultimately dependent on the user’s phenotype of interest. For example, isolating cells based on their positional phenotype within the population can be captured at a lower resolution. Isolating cells based on organelle level distinctions (such as differences in mitochondria localization) can require higher resolution. **? TROUBLESHOOTING**
36. Configure laser parameters. **! CAUTION** The two options for configuring multichannel imaging are (i) simultaneous or (ii) sequential scanning. Simultaneous scanning images every channel at the same time. Sequential scanning images each channel independently and can switch after every line, frame, or stack. A disadvantage to simultaneous scanning is that overlap in the emission spectra of two dyes will lead to crosstalk and caution should be taken. The Leica LAS X software provides scanning recommendations and can be referenced during set-up.

(a) Open the laser configuration window for the 405 nm, 488 nm, and 561 nm laser lines.
(b) Set the 488 nm and 561 nm laser lines to sequential scanning to avoid emission spectra crosstalk. Use the ‘between line’ scanning feature so live cell movement does not lead to signal artifact between channels. **CRITICAL** Short pixel dwelling time during speed scanning (800 Hz) to visualize Dendra2-green localization with 488 nm excitation light, likely will not cause Dendra2 photoconversion.
(c) Visualize the sample and adjust the objective to the desired focal plane.
(d) Use live view to adjust the detector, gain, and laser intensities for optimal exposure. **! CAUTION** Maintain minimal laser power output for visualizing Dendra2-green and keep in mind that different microscopes will vary in required laser power intensity; however, in all cases too high laser power will result in photobleaching, cell toxicity, and cell death. Phototoxicity can be assessed utilizing common cell viability assays, like Annexin V staining (Step 54).
(e) Use the photomultiplier tube (PMT) detector for photoconversion because during the bleach sequence, high intensity illumination targets the region of interest (ROI). The hybrid detector (HyDs) may switch off to protect the ROI from photon overload, resulting in loss of the post-bleach sequence.

### Photoconversion

**Timing** 1 - 4 h (photoconversion of user-defined and phenotypically distinct cells)

37. The photoconversion parameters set-up utilizing the FRAP interface. **CRITICAL STEP** Most confocal microscope software is equipped with a Fluorescence Recovery After Photobleaching (FRAP) feature. The FRAP parameters are then suitable for photoconversion.

(a) Turn the zoom-in feature on to allow for precise illumination and ROI selection.
(b) Set the background to zero to ensure that the area outside of the ROI is not exposed to background light. This will decrease the potential for false positive cell photoconversion near the ROI.
(c) Set the 405 nm laser intensity for photoconversion. This is dependent on cell type and experimental conditions. It is important to test various intensities to determine which intensity maintains cell viability while ensuring complete photoconversion for cells within the ROI (Box 2, Fig. 5B – D). **? TROUBLESHOOTING** **! CAUTION** Photobleaching occurs when laser intensity for photoconversion is too high and permanently eliminates fluorescent signal within the ROI due to a photon-induced covalent modification. If photobleaching occurs, decrease the 405 nm laser line intensity, and select a new ROI.
(d) Establish a time course for the number of prebleach, bleach, and post-bleach frames. These values are dependent on the experimental design. Optimize the time course by testing various conditions for each sample in study (Box 2). During these screening stages, monitor cells for any unhealthy signs, such as swelling or shrinkage. **? TROUBLESHOOTING**
38. Select the phenotype-driven ROI and begin the established time course for photoconversion.
39. Repeat steps 37 – 38 as necessary until all user-defined cells are photoconverted.
40. Assess efficiency of photoconversion within the evaluation menu on the user interface (Fig. 4B, C). The integrated density relative fluorescence values pre-and post-photoconversion are shown. The Dendra2-green should significantly decrease and Dendra2-red should significantly increase after photoconversion (Fig. 4B – E). **CRITICAL** The H2B-Dendra2 offers more accurate results over pal-Dendra2 because the 405 nm laser line can photoconvert within a singular region (versus around the periphery of the cell) increasing localized excitation and decreasing off-target photoconversion.

### Fluorescence-activated cell sorting (FACS)

**CRITICAL** Due to the different culture conditions that can be used with the SaGA platform, we discuss options for preparing non-adherent cells and samples cultured in 2D and 3D conditions for FACS.

### Option A: Non-adherent sample preparation for FACS

**Timing** 0.5 h (cell collection)

41. Centrifuge non-adherent cells at 125 – 350 g RT for 5 min. Centrifuge speeds may vary by cell line.
42. Proceed to step 51.

### Option B: 3D cell culture sample preparation for FACS

**Timing** 1.5 h (matrix degradation), 0.5 h (spheroid dissociation)

43. Dilute stock Collagenase/Dispase (C/D) cocktail in sterile media without serum for a working concentration between 1 – 5 mg/mL. For digestion of rBM 1 mg/mL, is sufficient. For digestion of type I collagen, a higher concentration is recommended. Enzyme concentration can be increased if the matrix is difficult to degrade.
44. Option 1: Mince matrix into quarters and place into a microcentrifuge tube with 3 – 4X volume of the minced matrix with the working stock of C/D. Place in a 37 °C incubator. Lightly vortex every 5 – 10 min until matrix is digested.
45. Option 2: Digest the matrix directly in the glass bottom dish. Remove all media prior to adding 3 – 4X volume of the working stock of the C/D digestion buffer. Place dish in a 37 °C incubator. Pipette gently every 5 – 10 min until matrix is digested and cells are released. **? TROUBLESHOOTING**
46. Centrifuge cells at 150 – 300 g RT for 5 – 10 min
47. Resuspend in trypsin (or similar proteolytic enzyme suitable for cleaving cell – cell junctions) to further digest spheroids and cell clusters into single cells. A standard light microscope can be used to visualize the formation of single cells. **? TROUBLESHOOTING**
48. Inactivate trypsin with TNS at a 1:1 ratio and centrifuge cells at 125 – 250 g RT for 5 min. Centrifuge speed may vary depending on cell line.
49. Proceed to step 51.

### Option C: 2D cell culture sample preparation for FACS

**Timing** 1 h (cell collection), 0.5 h (live/dead staining), 0.5 h (FACS cellular preparation)

50. Repeat steps 12.a.i – 12.a.vi for cell collection
51. Wash cells 1X by resuspending the cell pellet in 5 mL of flow buffer.
52. Centrifuge sample for 5 min at 250 g RT to pellet cells.
53. Gently resuspend approximately 1.0 × 10^6^ cells or less into 100 μL flow buffer. For samples with more than 1.0 × 10^6^ cells, resuspend in a larger volume to avoid cell aggregation or flow cytometer clogging.
54. Add 5 μL per 100 μL cell suspension of Annexin V conjugate (per manufacturer recommendations) and incubate for 10 min at RT. Protect samples from light from this point forward. **CRITICAL STEP** Annexin V conjugate will bind to phosphatidylserine, a marker of an apoptotic cell when exposed on the outer leaflet of the plasma membrane. Marking dead cells in the sample will reduce autofluorescence and increase population resolution to accurately select living, photoconverted cells.
55. Add 200 μL flow buffer per 100 μL solution. Gently pipette to mix and transfer to a 5 mL flow tube with a cell strainer. Place sample on ice for transport to the FACS sorter.
56. Prepare a tube with 1 mL of complete culture medium for collection of cells after FACS. Prior to sorting, run each sample through cell strainer flow tube cap to decrease cell aggregation.

### FACS set up

**Timing** 1 h (instrument set-up and sterilization)

57. Ensure waste tank is empty and sheath tank is at least 50 % full. Sheath tank should be tightly closed to allow for pressure build up in the system.
58. Power on the computer and activate the compressed airline to at least 80 psi.
59. Switch on the cell sorter and start the sorting software, ensuring the computer has successfully connected to the FACS sorter.
60. Initiate the fluidics start-up and follow the popup tutorial to complete required steps. This start-up will take approximately 10 min.
61. Remove the closing nozzle from the flow cell assembly and attach the appropriate nozzle for your experimentation. **CRITICAL STEP** Nozzle size is dependent upon cell size. For example, cells that are greater than 25 μm are best suited for a nozzle that is 100 μm or greater in size. Ensure that the software configuration matches the nozzle size of choice.
62. Activate the stream and stabilize to desired flow rate.
63. Calibrate the cytometer with cytometer set-up software and tracking beads. Set drop delay using Accudrop beads. This calibration will take approximately 20 min.
64. Sterilize the cytometer interior with 70 % (vol/vol) EtOH.
65. Sterilize the sample line with 10 % (vol/vol) bleach and rinse with water.

### FACS collection

**Timing** 1 h (evaluating control samples and isolating photoconverted cells)

66. Use lasers blue (B) 488 nm, yellow/green (YG) 561 nm, red (R) 633 nm, to excite and visualize the following, respectively: Dendra2-green – bandpass filter: B 530/30, Dendra2-red – bandpass filter: YG 582/15, Annexin V (Alexa Fluor 680) – bandpass filter: R 710/50. Additionally, Annexin V pacific blue (laser line: violet 405 nm, bandpass filter: 450/50) can be used in conjugation with Dendra2. Due its low intensity, the 405 nm and 488 nm lasers will not photoconvert Dendra2 while in the FACS sorter.
67. Prepare a workbook to collect live, single cells.
68. Perform the respective controls (as described in experimental design, Fig. 5A) and then flow sort cells positive for Dendra2-red into the collection tube prepared in Step 56 (Fig. 5A). These cells are the phenotypically driven, user-defined cell subpopulation of interest. **? TROUBLESHOOTING**
69. Process, culture, or store the collected cells based on the analysis of interest (Table 1, Fig. 2, Fig. 6).

### Downstream analysis post-FACS

Timing N/A (dependent upon assay of choice)

70. To sort cells for immediate bulk -omic analysis, collect the same number of cells for each photoconverted population into the desired lysis buffer (such as DNA or RNA lysis buffer) (Table 1).
71. To sort for immediate single cell -omic analysis, sort single photoconverted cells into individual wells of a 96-well plate with desired lysis buffer and immediately place in a negative 80 °C freezer (Table 1).
72. For cell propagation and long-term phenotypic analysis, sort cells into complete growth medium (Table 1). Culture and passage the cells as routinely done for further experimentation. **? TROUBLESHOOTING**

### TIMING

Steps 1 – 12, thaw and maintenance of tumor cells: 1 h thaw cells, 1 week tumor cell growth and maintenance

Steps 13 – 29, non-adherent, 3D, or 2D sample preparation for photoconversion: 1 h cell collection, counting, and plating (non-adherent and 2D) 1 h cell plating, 72 h incubation, 2 h embedding into matrix, 24 – 48 h incubation for invasion overtime (3D)

Steps 30 – 36, microscope and laser set up: 2 h incubator equilibration, 0.5 h microscope set up

Steps 37 – 40, photoconversion: 1 – 4 h photoconversion of user-defined and phenotypically distinct cells

Steps 41 – 56, non-adherent, 3D, or 2D sample preparation for fluorescence activated cell sorting: 0.5 h non-adherent cell collection, 1.5 h 3D matrix degradation, 0.5 h spheroid dissociation, 1 h 2D cell collection, 0.5 h cellular live/dead staining, 0.5 h FACS cellular preparation

Steps 57 – 65, FACS set up: 1 h instrument set-up and sterilization

Steps 66 – 69, FACS collection: 1 h evaluating control samples and isolating photoconverted cells

Steps 70 – 72, downstream analysis post FACS: N/A dependent upon assay of choice

### TROUBLESHOOTING

Troubleshooting advice can be found in Table 3.

**Table 3.** I Troubleshooting table

| Step | Problem | Possible reason | Solution |
| --- | --- | --- | --- |
| 18 | Cells are unable to form spheroid | Low cell - cell adherence junction expression, low incubation time | Repeat centrifugation (step 19) and/or incubate for an additional 24 h. |
| 33 | Cells are shrinking; detaching from plate; swelling | Inadequate cell culture conditions on tabletop incubator | Ensure that the incubator is working at the appropriate temperature, pressure, and CO<sub>2</sub> level. |
| 35 | Poor imaging resolution | Scanning pixel size and/or line averaging amount is too low | Increase these imaging acquisition parameters to increase resolution. |
| 37c | No fluorescent signal in the red channel | Laser intensity value is too low resulting in low to no photoconversion | Increase number of repetitions and/or bleach iterations. May need to increase laser intensity. |
| 37c | No fluorescent signal in the red channel | Laser intensity value is too high resulting in photobleaching ROI | Disregard ROI, decrease laser intensity, and select another ROI. |
| 37c | Cell shrinking or swelling after photoconversion | Laser intensity too high resulting in phototoxicity | Disregard ROI, decrease laser intensity, and select another ROI. |
| 37d | Low fluorescence in the red channel | Low photoconversion efficiency | Increase number of repetitions and/or bleach iterations. May need to increase laser intensity. |
| 44,45 | Unable to degrade matrix | Enzyme concentration too low, inadequate incubation time | Increase enzyme concentration or incubation time. Agitate matrix with pipette tip more frequently to encourage degradation. |
| 47 | Unable to degrade cell-cell junctions within spheroid | Enzyme volume or concentration too low, inadequate incubation time | Increase concentration, volume, or incubation time. Gently vortex to encourage junction cleavage. |
| 68 | Number of cells recovered is higher than number of photoconverted cells | Off-target photoconversion due to inadequate ROI placement and/or autofluorescence | Create a stricter ROI to ensure no off target or false positive photoconversion of nearby cells. Some cell types emit autofluorescence, ensure cytometer voltage settings are set to allow for enough separation between those autofluorescent cells and those that were photoconverted. Decrease laser intensity or time course on microscope to reduce off target photoconversion. |
| 68 | Low cell viability post FACS | Inadequate sample preparation and/or maintenance | Keep cells on ice to slow intracellular metabolism and increase survival. Avoid generating a dry pellet or air bubbles during processing. Air bubbles may create a surface tension that is toxic to the cells. Avoid vigorous vortexing and instead mix with gentle pipetting. If cell centrifugation is necessary post FACS, apply low speeds (125 - 250 g RT). |
| 72 | Poor cell proliferation and propagation | Poor collection conditions; not enough cells; crucial growth factors not present | Sort into culture media with at least 20% FBS to increase growth factors and promote cell survival. Coat cultivation plates with protein to promote cell adhesion. Plate cells on smaller surface area plate to facilitate cell - cell communication to promote cell survival. |

## ANTICIPATED RESULTS

The SaGA platform affords the unique ability to isolate live cells based upon image-able whole cell or organelle morphological distinctions. The initial technical setup for SaGA, such as stable introduction of photoconvertible tags or determining imaging parameters, may require initial troubleshooting. However, once established, live cell spatiotemporal multi-omic analysis can be performed utilizing the same experimental parameters.

### Cell transfection and transduction

Dendra2 can be engineered onto a variety of protein targets. Importantly, introducing exogeneous tags to a protein can result in altered protein function and/or activity, and potentially feed forward within the experimental system resulting in cellular, population and subpopulation behavioral changes. To this end, with both the H2B-Dendra2 and pal-Dendra2 PCFPs, we experimentally confirmed that our cellular attributes and phenotypes of interest (proliferation, cell-cell junctional integrity, biomarker expression, spheroid formation, collective invasion) were maintained upon addition of Dendra2 (Fig. 3). Similarly, when applying the SaGA platform to alternate research questions, we recommend performing similar experimental comparisons to probe the phenotype of interest with and without the protein tag.

### Live cell imaging and photoconversion

Isolating viable cells based upon a phenotype of interest requires defining an experimental window in which the phenotype occurs. For example, after monitoring H1299 3D spheroid invasion, we determined that day 5 presented clear leaders and followers within the collective invasion pack that can be photoconverted (Fig. 2). This time course will vary depending on experimental design. Most scanning confocal microscopes are equipped with scanning-FRAP modes that can be adapted to photoconvert live cells within a specific region of interest.

### Fluorescence activated cell sorting

FACS is optimally performed when at least 50 cells or greater are photoconverted for sorting to account for cell loss during the cell preparation, selection, and isolation (Steps 39 – 56). The inclusion of a live/dead stain aids in the sorting of viable cells and to assess any death due to laser intensity or 3D degradation mechanisms. To enhance the number of live photoconverted cells during FACS, we recruited colleagues to streamline isolation steps. One person was designated to either photoconvert (Steps 30 – 40), prepare cells for FACS (Steps 41 – 56), or sort cells for analysis (Steps 57 – 69). Utilizing this methodology, we were able to photoconvert and sort 100s of live cells in one work day for downstream analysis.

### Downstream application and analysis example

Utilizing SaGA-derived leader and follower cells, we performed multi-omic analysis to extrapolate DNA methylation status and bulk transcriptomic profiling^26,28–30^. We established that leaders and followers maintain stable differences at the epigenetic, genetic, metabolomic, and transcriptomic level^26–30^. Similarly, propagation of isolated leader and follower subpopulations generate stable phenotypes for over 30 passages (Fig. 2F)^26^. Together, these data showcase the molecular significance in phenotypic positioning.

In this protocol, we apply SaGA to define transcriptomic differences within leaders and followers at the single cell level by isolating cells actively undergoing 3D collective invasion in recombinant basement membrane. These cells were immediately sorted into 96-well plates including RNA lysis buffer for downstream single cell RNA-sequencing (Fig. 6A). After library preparation and next-generation sequencing, samples were filtered by library size, number of expressed features, and proportion of mitochondrial reads, resulting in 105 total cells. The top 6% most variably expressed genes (i.e. the core gene set; n=1,155 genes) across all single cells were defined by a combined variance, median absolute deviation (MAD) and IQR statistic. Consensus clustering was performed on the core gene set using Canberra distance, complete clustering with 1000 iterations, 80 % sample resampling and 100 % gene resampling. A tSNE plot was generated for the core cell set based upon the core gene set (Fig. 6B). Previously established leader-and follower-specific gene mutations were used to complement positional phenotypes to label each single cell, enabling for the first time, the combination of phenotypic, transcriptomic, and mutational analysis of single cells during active collective invasion^29,30^. We selected mutation loci from the list of leader-and follower-specific genes and verified that they had sufficient coverage in the single-cell sequences. This left 5 leader-specific mutation loci, and 7 follower-specific mutation loci (Fig. 6C). Interestingly, there was 95.9 % mutual exclusivity between the mutational profiles on the single-cell level – that is, of the 105 cells with at least one leader-or follower-specific mutation, only 7 cells (4.1 %) had both (Fig. 6D). Next, the correlation between mutation profile and specific leader-follower gene expression markers was measured. To determine the genes driving the formation of two separate leader and follower clusters in the gene expression tSNE plot, we performed gene set enrichment analysis (GSEA) for two groups: one comprised of tSNE clusters 1 and 2, and the other comprised of tSNE clusters 3 and 4. Among the mostly highly enriched gene sets in tSNE clusters 1 and 2 were cell cycle and cell proliferation, which included core enrichment of genes including CCNA2/B1/B2, CDK1, and Ki-67, suggesting that tSNE clusters 1 and 2 represented more actively proliferating populations. By contrast, tSNE clusters 3 and 4 were not enriched for cell cycle or proliferation gene sets. Among the most differentially expressed leader cell markers from our previous bulk RNA-sequencing was MYO10, an unconventional myosin involved in filopodial elongation^28^. Upon analyzing the expression of MYO10 within each single cell comprising the four tSNE clusters, we found that Clusters 1 and 4 had significantly increased expression of MYO10 compared to clusters 2 and 3 (Fig. 6E-F). These results highlight the valuable molecular information underlying live-cell spatial organization. We envision that SaGA can be applied to a broad range of experimental studies to further exploit distinct cellular responses within a parental population, thereby continuing to identify critical cell subpopulations and their mechanistic dependencies.

## Author contributions

A.I.M. and J.M.K. conceived the original SaGA technique; T.O.K., A.M.A. and J.K.M. contributed to the expansion and further development of SaGA; T.O.K. and A.M.A. drafted and revised the manuscript; T.O.K., A.M.A., and B.P. designed the figures; A.M.A., J.K.M., T.O.K., E.R.S., N.M.Z., and B.P. carried out the experiments; A.I.M., T.O.K, A.M.A. and B.P. contributed to the interpretation of the results; J.K.M., C.M.K., and A.I.M. edited the manuscript; A.I.M. obtained funding for this project; T.O.K. managed the manuscript preparation; all authors read, edited, and approved the final manuscript.

